# Bin-Free Fitting of Probability Distributions Emphasizing DNA Histograms

**DOI:** 10.1101/218933

**Authors:** Nash Rochman

## Abstract

It is often challenging to find the right bin size when constructing a histogram to represent a noisy experimental data set. This problem is frequently faced when assessing whether a cell synchronization experiment was successful or not. In this case the goal is to determine whether the DNA content is best represented by a unimodal, indicating successful synchronization, or bimodal, indicating unsuccessful synchronization, distribution. This choice of bin size can greatly affect the interpretation of the results; however, it can be avoided by fitting the data to a cumulative distribution function (CDF). Fitting data to a CDF removes the need for bin size selection. The sorted data can also be used to reconstruct an approximate probability density function (PDF) without selecting a bin size. A simple CDF-based approach is presented and the benefits and drawbacks relative to usual methods are discussed.

## I. INTRODUCTION

Representing a noisy experimental data set can be challenging[3], and searching for the right, bin size to construct a histogram is a common problem. For example, it is often a struggle to assess whether a cell cycle synchronization (in G1) experiment was successful or not. In this case, the goal is to determine whether the DNA content (as inferred from a Hoechst stain, DAPI stain. etc.)[1][4] is best represented by a unimodal distribution indicating that the majority of the population has a single copy of DNA, and the synchronization was successful, or that the ensemble is better represented by a bimodal distribution, and the synchronization was unsuccessful [2]. An example of the asynchronous case is displayed in Fig. 1.

**Figure 1:**
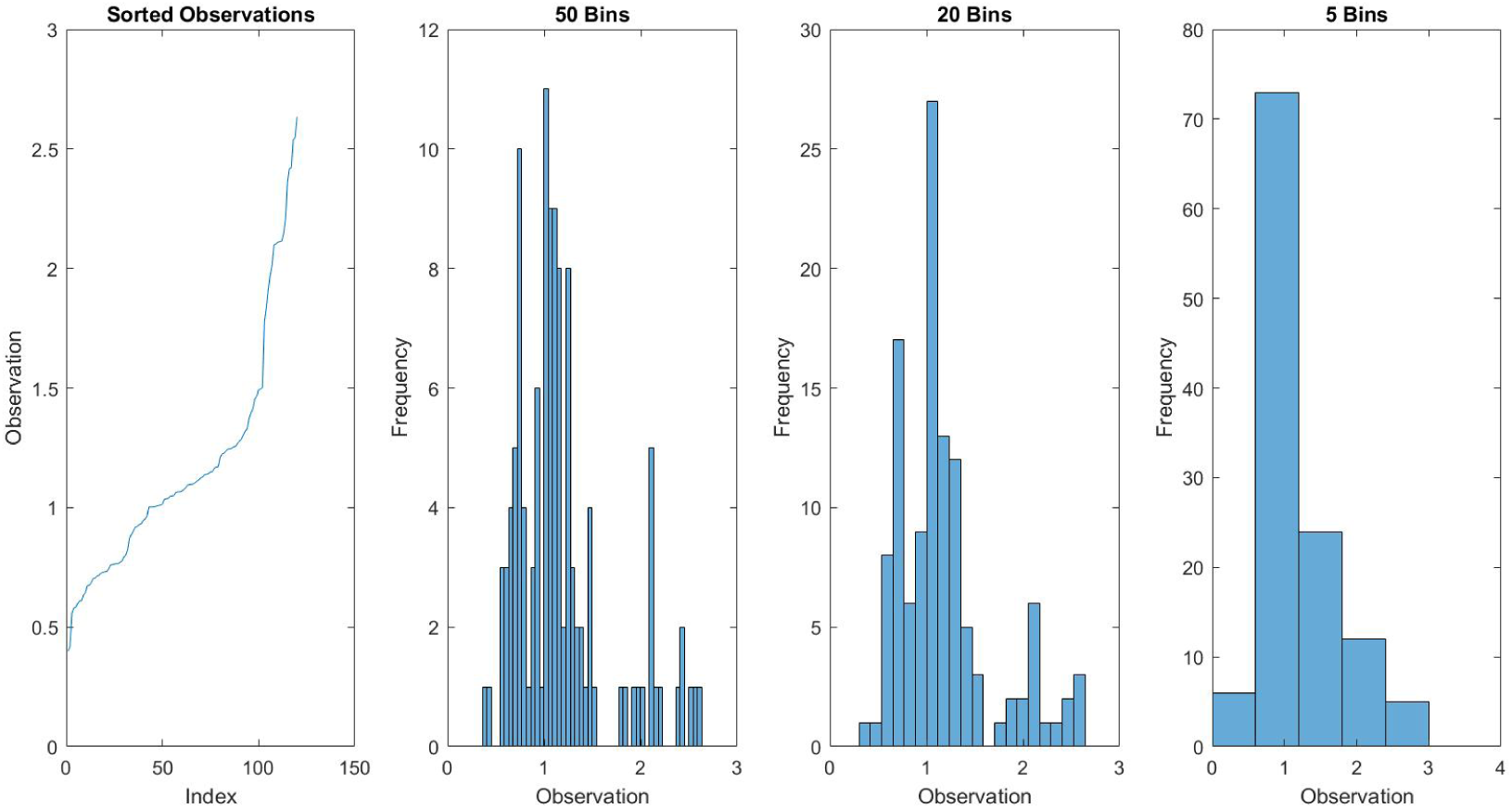
An illustration of the problem: the goal is to determine if the ensemble from which these observations (e.g. single cell DNA content) are sampled is unimodal or bimodal.

This simulated data was sampled from a bimodal distribution, 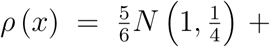 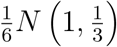 where N (*μ, σ*) is a normal distribution of mean *μ* and standard deviation *σ*; but when divided into fifty bins, one might, conclude it, is simply right, skewed. Similarly looking at, five bins, the distribution appears unimodal. The selection of twenty bins more convincingly displays the bimodality; however, the underlying issue remains the same: the selection of bin size is subjective. The bin size is selected to approximate the probability density function (PDF) from which the data was drawn. Data are often described by their PDFs because many basic properties (mean, mode, etc.) can be conveyed easily this way. This is inconvenient when it comes to fitting a functional form, however, because it requires both picking a bin size and also normalizing the resultant distribution. Alternatively, the data can be fit to the cumulative distribution function without these steps.

## II. FITTING TO THE CDF

Fitting to the cumulative distribution function, 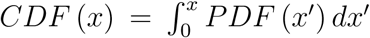, directly avoids the need for selecting a bin size. Consider *N* data points. Sorted, from smallest to largest, point *n_i_* represents a measurement of the 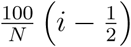 percentile of the distribution. We can calculate these percentiles for a test function by finding the points at which the CDF is equal to 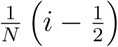:

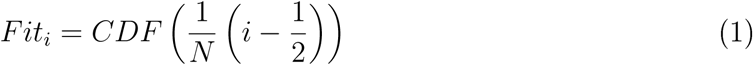

It is then very straightforward to optimize the test function - simply choose parameters which minimize the least squared error of the *N* fit points and the date. See Fig. 2.

**Figure 2:**
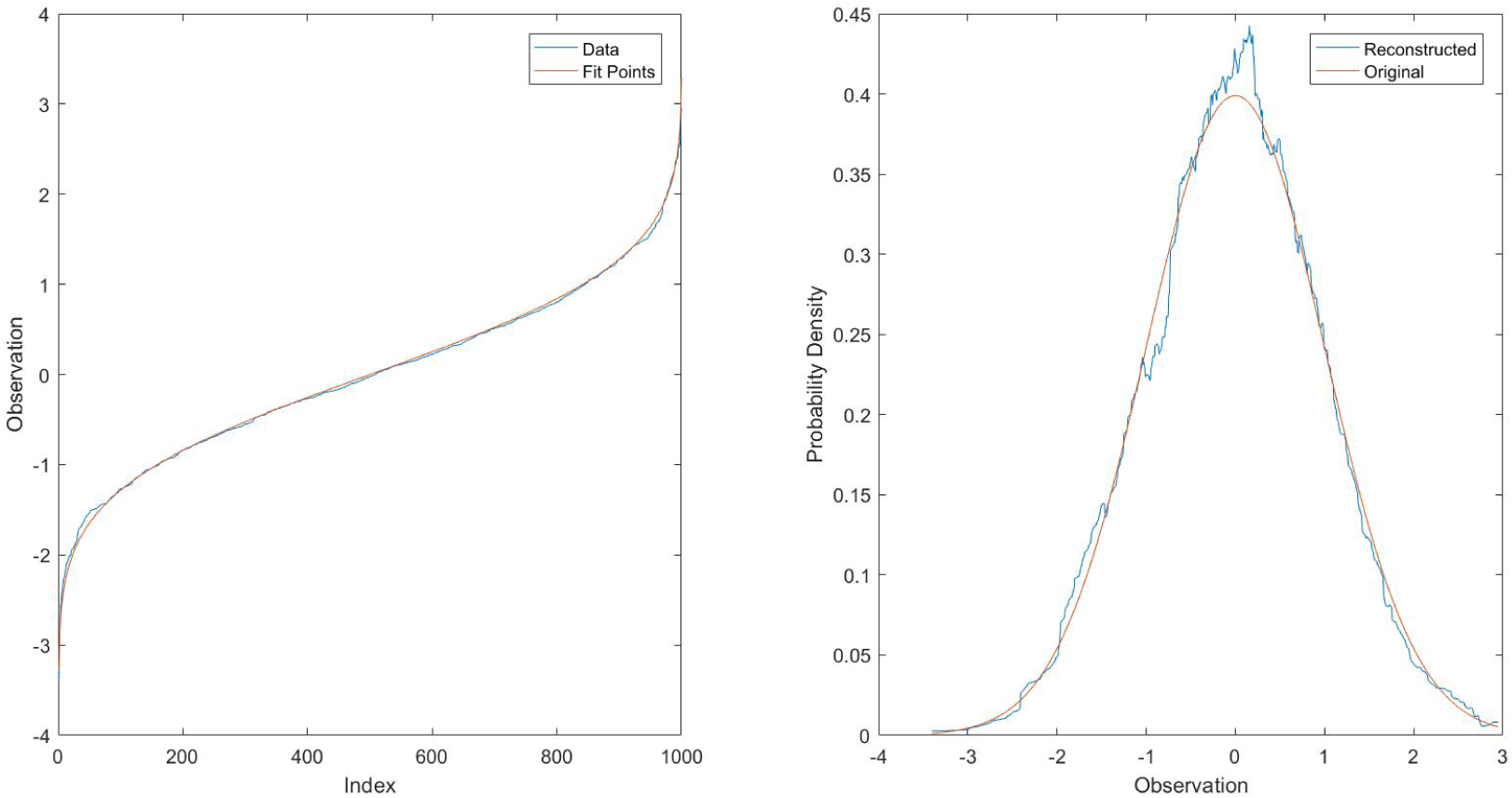
An illustration of the CDF based method. The data is drawn from *N* (0,1). On the left is a comparison between the sorted data and the points 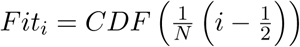. On the right is the probability density function reconstructed from the sorted points and smoothed with a moving average over 150 points.

On the other hand, if an estimate of the probability density function is desired, an approximation can be constructed from the the sorted points. The probability density from point 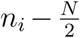 to 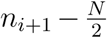 may be approximated to be 1/(*N*(*n_i+i_* − *n_i_*)) since the probability an observation will fall between these points is assumed to be 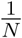 and the distance between the points is just *n_i_*_+1_ – *n_i_.* The reconstructed distribution is very noisy and a smoothing of this construction may have greater utility; however, it should be noted that this smoothing is just, as arbitrary as the bin size selection.

## III. DISCUSSION

Fitting sorted data to a cumulative distribution function of a suggested functional form avoids the selection of bin size which can introduce unwanted subjectivity. The simple method above works well when a model functional form is known or desired. See Fig 3.

**Figure 3:**
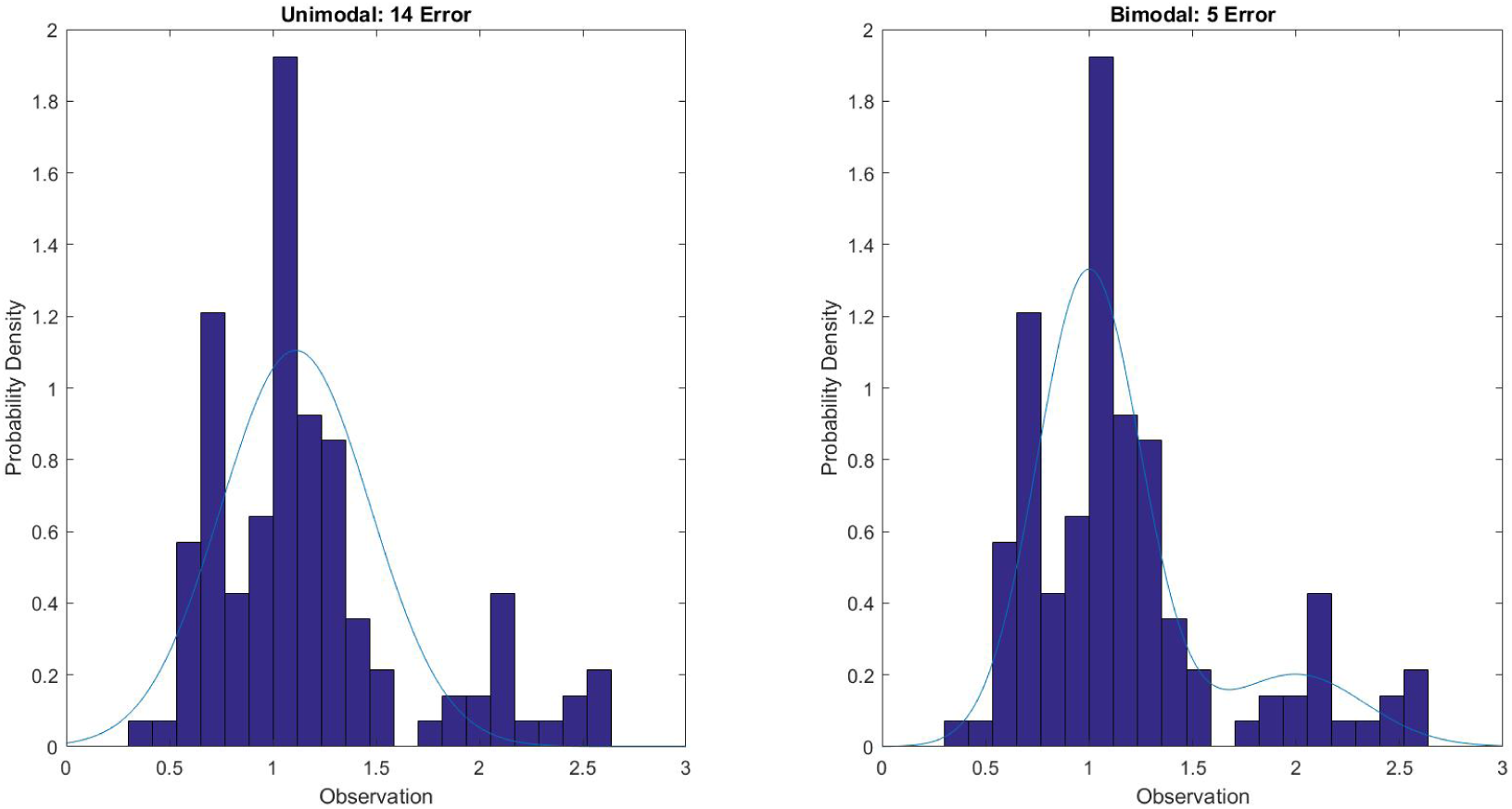
A demonstration of the improved fit from the CDF based method for a bimodal distribution. The best fit unimodal, normal distribution is *N* (1.1, 0.36). The bimodal distribution tested is the parent distribution 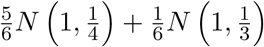.

With the CDF based method, different model functions can easily be tested against one another (e.g. a unimodal Gaussian or bimodal Gaussian sum as displayed in Fig. 3) without any qualitative assessment of the PDF. This may be useful for new students or anyone who is uncertain about how to pick a “good” bin size for PDF approximation.

